# *Clostridioides difficile* colonization is not mediated by bile salts and utilizes Stickland fermentation of proline in an *in vitro* model

**DOI:** 10.1101/2024.07.17.603937

**Authors:** Xiaoyun Huang, April E. Johnson, Joshua N. Brehm, Thi Van Thanh Do, Thomas A. Auchtung, Hugh C. McCullough, Armando I. Lerma, Sigmund J. Haidacher, Kathleen M. Hoch, Thomas D. Horvath, Joseph A. Sorg, Anthony M. Haag, Jennifer M. Auchtung

## Abstract

Treatment with antibiotics is a major risk factor for *Clostridioides difficile* infection, likely due to depletion of the gastrointestinal microbiota. Two microbiota-mediated mechanisms thought to limit *C. difficile* colonization include conversion of conjugated primary bile salts into secondary bile salts toxic to *C. difficile* growth, and competition between the microbiota and *C. difficile* for limiting nutrients. Using a continuous flow model that simulates the nutrient conditions of the distal colon, we investigated how treatment with six clinically-used antibiotics influenced susceptibility to *C. difficile* infection in 12 different microbial communities cultivated from healthy individuals. Antibiotic treatment reduced microbial richness; disruption varied by antibiotic class and microbiota composition, but did not correlate with *C. difficile* susceptibility. Antibiotic treatment also disrupted microbial bile salt metabolism, increasing levels of the primary bile salt, cholate. However, changes in bile salt did not correlate with increased *C. difficile* susceptibility. Further, bile salts were not required to inhibit *C. difficile* colonization. We tested whether amino acid fermentation contributed to persistence of *C. difficile* in antibiotic-treated communities. *C. difficile* mutants unable to use proline as an electron acceptor in Stickland fermentation due to disruption of proline reductase (*prdB-*) had significantly lower levels of colonization than wild-type strains in four of six antibiotic-treated communities tested. Inability to ferment glycine or leucine as electron acceptors, however, was not sufficient to limit colonization in any communities. This data provides further support for the importance of bile salt-independent mechanisms in regulating colonization of *C. difficile*.

**IMPORTANCE:** *C. difficile* is one of the leading causes of hospital-acquired infections and antibiotic-associated diarrhea. Several potential mechanisms through which the microbiota can limit *C. difficile* infection have been identified and are potential targets for new therapeutics. However, it is unclear which mechanisms of *C. difficile* inhibition represent the best targets for development of new therapeutics. These studies demonstrate that in a complex *in vitro* model of *C. difficile* infection, colonization resistance is independent of microbial bile salt metabolism. Instead, the ability of *C. difficile* to colonize is dependent upon its ability to metabolize proline, although proline-dependent colonization is context-dependent and is not observed in all disrupted communities. Altogether, these studies support the need for further work to understand how bile- independent mechanisms regulate *C. difficile* colonization.

## INTRODUCTION

*Clostridioides difficile* is one of the leading causes of nosocomial infections due to its transmissibility as environmentally-resistant spores and its ability to infect patients who have been treated with antibiotics (1, 2). Many classes of antibiotics can disrupt the colonic microbiota (3–5); disruption can decrease production of inhibitory metabolites (6–9) and reduce competition for limiting nutrients (10–13), providing favorable conditions for *C. difficile* infection. Although multiple mechanisms for colonization resistance have been identified, an understanding of the hierarchical importance of these mechanisms in *C. difficile* colonization and disease is just beginning to emerge.

Microbial bile salt metabolism has been extensively studied for its role in limiting *C. difficile* colonization. The primary bile salts cholate and chenodeoxycholate are synthesized in the liver and are conjugated to glycine or taurine to improve solubility (14). Once secreted into the intestine, microbial enzymes begin modifying these bile salts, removing conjugated amino acids and dehydroxylating primary bile salts into secondary bile salts (14). *C. difficile* spore germination is stimulated by cholate-family bile salts (cholate, taurocholate, glycocholate, deoxycholate) and inhibited by chenodeoxycholate-family bile salts (15, 16), although there is variation in these responses between strains (17). Germination is enhanced by amino acid (15, 18) and calcium co-germinants, which act synergistically with bile salts to enhance germination (19). Secondary bile salts (deoxycholate, lithocholate) inhibit growth of vegetative *C. difficile* in vitro (7, 15, 20), and low levels of secondary bile salts correlate with *C. difficile* infection in humans (21, 22) and in mouse models (7, 23, 24).

However, recent studies have demonstrated that our understanding of the role of bile salts in *C. difficile* colonization resistance may be incomplete. *C. difficile* spore colonization and fulminant disease was observed in *Cypb8b1*^-/-^ mice unable to make cholate-family bile salts (25), and resistance to *C. difficile* colonization was observed even though these mice do not produce the secondary bile salts deoxycholate or lithocholate. *Clostridium scindens*, a microbe inferred to inhibit *C. difficile* growth through dehydroxylation of primary bile salts to secondary bile salts (24), was also shown to inhibit *C. difficile in vitro* through production of tryptophan-derived antibiotics (9) and other uncharacterized metabolites (26) and to potentially compete with *C. difficile* for nutrients required for growth in *Cypb8b1*^-/-^ monocolonized mice (25). In addition, both cholate and deoxycholate were shown to induce similar *C. difficile* stress responses, although 10X higher concentrations of cholate were used to observe these effects (27).

Competition for nutrients between commensal microbes and *C. difficile* has long been postulated as a mechanism for colonization resistance, with Wilson and colleagues utilizing continuous-flow culture systems to demonstrate competition between the microbiota and *C. difficile in vitro* (28, 29). *C. difficile* can metabolize mucin monosaccharides *in vitro* (30), and preferentially expresses pathways for degradation of mucin monosaccharides in mouse models (31, 32), indicating that metabolism of mucin monosaccharides may be a niche open to *C. difficile* during infection. Similarly, *C. difficile* metabolism of carbohydrates and sugar alcohols from the host and its diet may be another niche open to *C. difficile* during infection, as expression of genes in these metabolic pathways increase during infection in germ-free and antibiotic-treated mice (11, 31, 32). In addition to carbohydrate metabolism, *C. difficile* also efficiently utilizes amino acids through Stickland fermentation (33), which is a metabolic pathway limited primarily to proteolytic Clostridial species (34), providing a unique metabolic niche within the GI tract. Increasing evidence from human (10, 35) and mouse (11, 13, 25, 31, 32, 36) studies points to Stickland fermentation with proline as an electron acceptor as a preferred nutritional niche for *C. difficile* in the GI tract. *C. difficile* can also use glycine (33) and leucine (37) as electron acceptors in Stickland fermentation, with a glycine reductase mutant recently shown to delay morbidity in a hamster model of disease (38). Multiple regulatory pathways converge to coordinately regulate the expression of proline, glycine, and leucine reductase pathways (34), and more work is needed to understand how the integration of these pathways contributes to disease in different environmental and nutritional contexts.

Previously, we described an *in vitro* minibioreactor array (MBRA) model for characterization of *C. difficile* colonization resistance in the presence of human fecal microbial communities under nutritional conditions that simulate the distal colon (39), that has subsequently been used to characterize microbes that inhibit *C. difficile* colonization and/or toxin production (40–42). In this study, we used the MBRA model to investigate how treatment with six clinically-used antibiotics differentially impacted susceptibility to *C. difficile* infection in microbial communities cultivated from 12 healthy individuals. As expected, we observed that antibiotic treatment reduced microbial richness and altered microbial diversity, although the extent of antibiotic-mediated disruption varied across individual communities and was not well correlated with susceptibility to *C. difficile* infection. Antibiotic treatment also reduced microbial metabolism of cholate to deoxycholate, although there was no correlation between deoxycholate levels and susceptibility to *C. difficile* infection. To further test whether bile salts contributed to *C. difficile* colonization resistance, we cultivated fecal communities in the presence and absence of bovine bile. Similar to what was observed by Aguirre et al. (25) in *Cypb8b1*^-/-^ mice, we observed bile salts were not required for *C. difficile* colonization resistance and that the low level of *C. difficile* spore germination that occurred in the absence of bile was sufficient to allow colonization of antibiotic-treated communities by *C. difficile* spores. Using *C. difficile* mutants unable to utilize proline, glycine, or leucine as electron acceptors during Stickland fermentation due to a mutation in proline reductase (*prdB* (43)) glycine reductase (*grdA*), or 2-hydroxyisocaproate dehydrogenase (*ldhA*), we demonstrated that proline reduction was required for *C. difficile* to colonize a subset of antibiotic-disrupted microbial communities, whereas glycine and leucine reduction was not required for *C. difficile* colonization in the same communities. These results can serve as a foundation for further mechanistic characterization of the hierarchical importance of different nutritional environments in limiting *C. difficile* colonization.

## RESULTS

### Loss of microbial richness is not sufficient to promote *C. difficile* colonization

Previously, we described an *in vitro* model for culturing microbial communities from fecal samples under conditions that simulated the nutritional environment of the distal colon (44). Communities were shown to maintain a subset of microbial richness and diversity found in the starting fecal samples, similar to other *in vitro* models (44) and germ-free animal models colonized with human feces (45, 46). Further, communities cultured from most healthy individuals demonstrated resistance to colonization by *C. difficile*, which could be disrupted by treatment with clindamycin (39, 41). To better understand how this model can be used to measure effects of antibiotics on microbiota disruption and *C. difficile* colonization, we compared treatment with six individual antibiotics (Fig. 1A) on microbial community composition and *C. difficile* susceptibility of communities cultured from twelve different healthy individuals. These antibiotics were selected because they are all used clinically, but vary in their therapeutic spectrum of antimicrobial activity.

**Figure 1:**
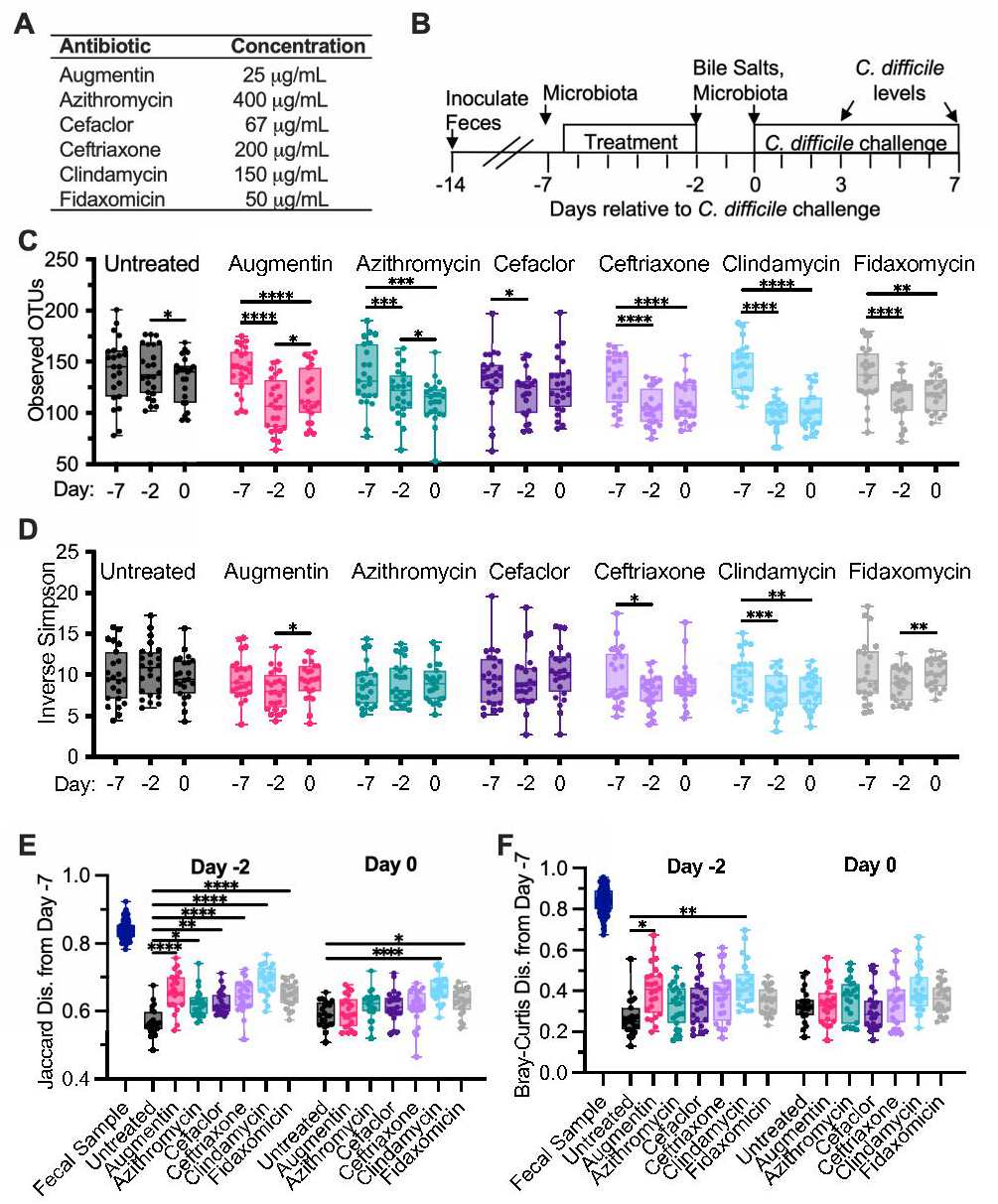
Alterations in microbiota diversity varied by type of antibiotic administered. (**A**) Antibiotics administered during experiment. (**B**) Experimental timeline indicating points where samples were collected; time is indicated relative to the point of *C. difficile* challenge (Day 0). In (**C**)-(**F**), samples were collected from duplicate reactors inoculated with one of twelve healthy human fecal samples and treated as indicated. (**C**) Microbiota richness (Observed OTUs with ≥ 99% identity) and (**D**) microbial diversity (Inverse Simpson measure) were determined from 16S rRNA gene sequences. Statistical significance of changes between time points (Days -7, -2, and 0) are indicated for each antibiotic. (**E**) Jaccard and (**F**) Bray-Curtis dissimilarities were determined from samples collected on Day -7 and plotted for each reactor and the fecal inocula. Statistical significance of differences between each antibiotic-treated sample and untreated communities at each time point are indicated. *, p<0.05; **, p<0.01; ***, p<0.001; ****, p<0.0001. All data is plotted, with boxes representing the interquartile range (IQR) and cross lines representing the median values.

Fourteen independent bioreactors were inoculated with each fecal sample and allowed to stabilize in continuous culture for seven days. Reactors were treated in duplicate with one of six antibiotics twice daily for five days (Figure 1B); two reactors served as untreated controls. Two days after the end of antibiotic treatment, communities were challenged with vegetative cells of *C. difficile* strain 2015 (CD2015), a ribotype 027 clinical isolate (39). Samples were collected for analysis of microbial community composition by 16S rRNA gene sequencing from three replicates of the initial fecal sample prior to inoculation, and from cultured communities prior to administration of antibiotics (Day -7), at the end of antibiotic treatment (Day -2), and two days later (Day 0), just prior to *C. difficile* challenge (Figure 1A). Samples were collected from cultured communities for analysis of *C. difficile* levels on days 3 and 7 post-infection.

Prior to antibiotic treatment (Day -7), there were no differences in richness (Observed OTUs with ≥99% average nucleotide identity, Figure S1A) or microbial diversity (Inverse Simpson, Figure S1B) between treatment groups, although there was a loss in richness and diversity compared to the fecal sample inocula as reported previously ((44); Figures S1A and S1B). Communities were composed primarily of taxa from the Bacteroidetes, Firmicutes, Proteobacteria, and Verrucomicrobia phyla, with members of Fusobacteria, Synergistetes, and Actinobacteria phyla found in some communities (Figure S2).

Treatment with antibiotics significantly reduced microbiota richness at the end of antibiotic treatment (Day -2) relative to the baseline sample collected from the same reactor at Day -7 (Figure 1C) and led to significantly lower levels of richness compared to untreated controls on Day -2 (Figure S1A); decreases ranged from 1.1-fold (cefaclor) to 1.5-fold (clindamycin). Microbial diversity declined significantly from baseline to the end of antibiotic treatment with just two antibiotics – clindamycin and ceftriaxone (Day -2, Figure 1D). These antibiotics also showed significant lower levels of diversity compared to untreated samples at Day -2, as did the Augmentin-treated communities (Figure S1B).

We also measured changes in shared community composition (Figure 1D) and structure (Figures 1E, F) by calculating Jaccard and Bray-Curtis dissimilarity measures from communities before antibiotic treatment (Day -7) to communities after antibiotic treatment (Day -2). (Jaccard and Bray-Curtis dissimilarity measures compare the proportion of taxa that are shared between two communities, with Jaccard providing an unweighted measure of dissimilarity between communities and Bray-Curtis providing a measure of dissimilarity that is weighted based on taxa abundance. In both cases, values closer to 0 are more similar.) As had been reported previously (44), cultivation led to significant shifts in microbial composition and structure from initial fecal samples (Figures 1E, F) to Day -7. Following this initial reorganization in community composition and structure, continued cultivation led to lower levels of change in community composition and structure (Figure 1E and 1F, see untreated samples on Day -2 and Day 0).

Treatment with all antibiotics led to larger changes in community composition from Day -7 to Day -2 compared to untreated communities (Figure 1E), whereas only Augmentin and clindamycin also led to significantly larger changes in community structure compared to untreated communities (Figure 1F).

Two days following the end of antibiotic treatment (Day 0), reduced richness compared to baseline samples persisted for all antibiotic-treated communities with the exception of those treated with cefaclor (Figure 1C), although Augmentin-treated communities exhibited small, but statistically significant increases in microbial richness (Figure 1C). Augmentin-treated communities also exhibited small, but statistically significant increases in microbial diversity (Figure 1D) and decreases in Jaccard (Figure S1E) and Bray-Curtis (Figure S1F) dissimilarity, indicating a potential return towards baseline following cessation of this antibiotic. Clindamycin-treated communities continued to exhibit decreased microbial diversity (Figure 1D) and less similar community composition to baseline (Figure 1E).

*C. difficile* susceptibility, measured as levels of *C. difficile* detected on Day 7 following challenge with *C. difficile* cells, varied across different antibiotic treatments (Figure 1E) and fecal donors (Figure S3). While *C. difficile* levels on Day 3 and 7 were collected, we used levels of *C. difficile* on Day 7 rather than Day 3 as a marker of colonization to provide sufficient time for loss of non-replicating cells and spores under continuous flow culture conditions. Of the twelve fecal samples tested, untreated communities from six fecal samples were resistant to colonization, defined as *C. difficile* colony forming units (CFU)/mL undetectable in both replicates, and untreated communities from one fecal sample were colonized at high levels, with both replicates colonized with *C. difficile* levels in the highest quartile. The remaining five untreated communities exhibited variable susceptibility to colonization (Figure S3A), with two fecal communities exhibiting low (1^st^ quartile) or undetectable levels of colonization in both replicates. Clindamycin was the only antibiotic that significantly increased median levels of *C. difficile* colonization above levels observed in untreated communities (Figure 2A); ten of twelve fecal sample communities were colonized following treatment with clindamycin (Figure S3A). Susceptibility to colonization varied across communities from different fecal samples that were treated with other antibiotics (Figure S3A).

**Figure 2:**
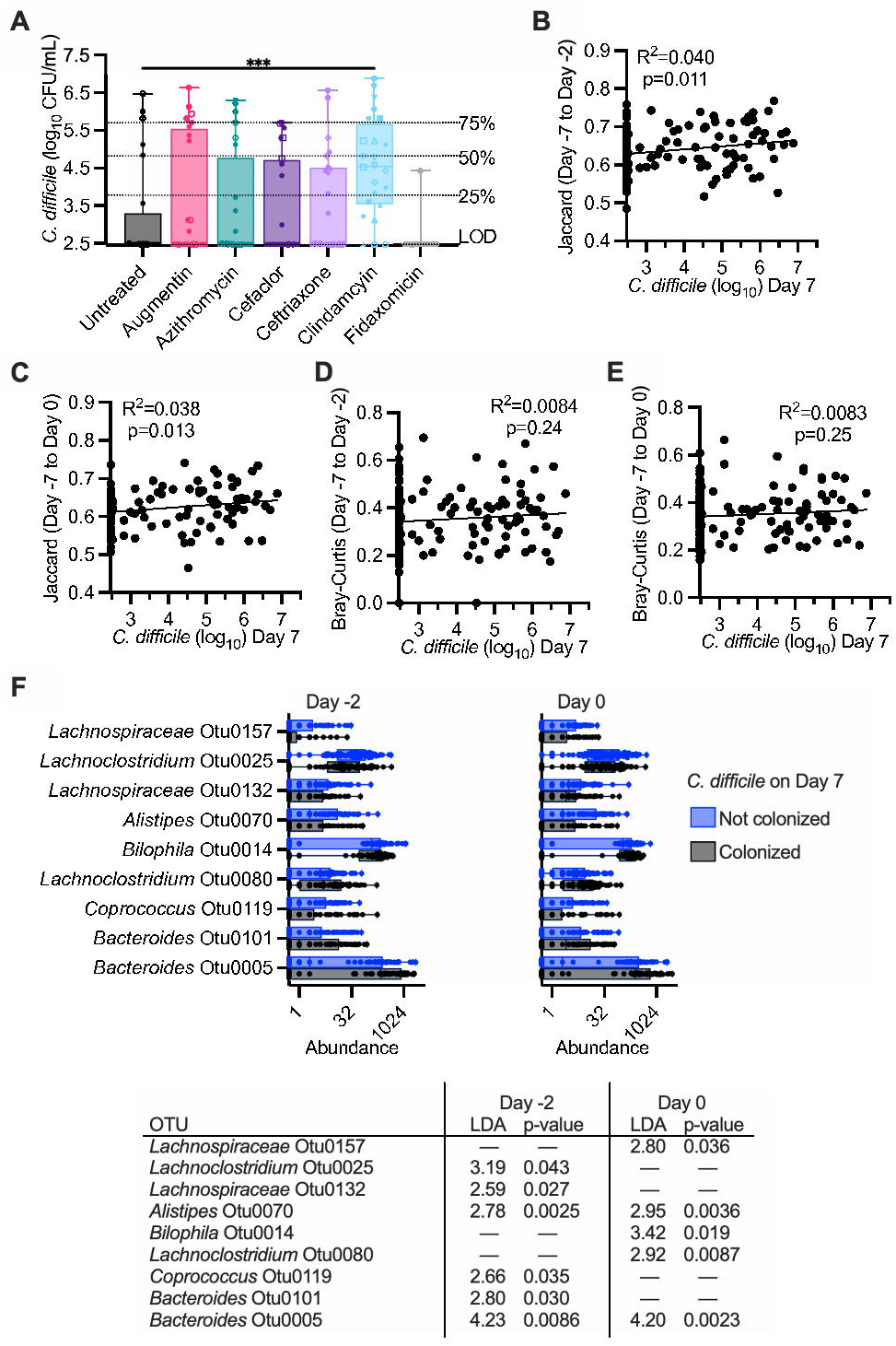
Clindamycin-treatment increases susceptibility to *C. difficile* colonization. **(A)** Levels of *C. difficile* measured in all reactors on Day 7 of infection. Data is pooled by treatment, with each reactor and replicate indicated by a distinct symbol consistent across treatment groups. (Effects on individual fecal samples can be better visualized in Supplementary Figure S3.) Statistical significance of differences between each antibiotic-treated community and untreated communities are indicated; ***, p<0.001. **(B)** – (**E)** Simple linear regression between levels of *C. difficile* on Day 7 and Jaccard (**B**-**C**) and Bray-Curtis (**D**-**E**) dissimilarity on Day -2 (**B**,**D**) or Day 0 (**C**,**E**). (**F**) OTUs that differed significantly in abundance on Day -2 and/or Day 0 in communities that were colonized or not colonized with *C. difficile* on Day 7 were identified with LEfSe analysis of rarefied OTU abundance data. Abundance of OTUs across all samples are plotted for colonized and not colonized communities, with linear discriminant analysis (LDA) and p-values determined by LEfSe reported below abundance plots.

We tested whether there were correlations between levels of richness or microbial diversity on Day -2 or Day 0 and found that there were no significant correlations with *C. difficile* colonization levels (Figure S4). We also assessed whether changes in community composition (Jaccard dissimilarity) or structure (Bray-Curtis dissimilarity) from Day -7 to Day -2 or Day 0 correlated with *C. difficile* colonization levels. We observed weak correlations with changes in microbial composition from baseline (Day -7) to Day -2 (Figure 2B) and Day 0 (Figure 2C). There were no significant correlations with changes in community structure from baseline to Day -2 or Day 0 (Figure 2E). Similar results were observed for *C. difficile* levels on Day 3 (Figure S5). We used LEfSe (47) to identify specific OTUs whose abundance on Day -2 or Day 0 correlated with *C. difficile* colonization. We found a small number of OTUs whose abundance on Day -2 or Day 0 significantly correlated with *C. difficile* levels (Figure 2F), with four OTUs more abundant in non-colonized communities on Day -2 and/or Day 0 and five OTUs more abundant in colonized communities on Day -2 and/or Day 0.

### Antibiotic treatment alters bile salt metabolism

Bioreactor medium (BRM3) contains bovine bile as a complex source of bile salts. Analysis of bioreactor medium indicated that cholate family bile salts predominated, with approximately 36% taurocholate (198 μM), 34% glycocholate (190 μM), 10% cholate (56 μM), and 2% deoxycholate (13 μM) of total bile salts measured (Figure S6A-B). To understand how microbiota cultivation and antibiotic treatment altered bile salt pools in bioreactor cultures, we focused on quantification of the amino acid conjugated primary bile salt, taurocholate, the primary bile salt, cholate, and the secondary bile salt, deoxycholate across the communities described in Figure 1. We observed that microbiota cultivation led to a >6,500-fold decrease in median levels of taurocholate and a 45-fold increase in median levels of deoxycholate in spent culture supernatant in untreated communities on Day - 2 compared to fresh medium (Figure S6C). Treatment with four antibiotics – Augmentin, ceftriaxone, clindamycin, and fidaxomicin – led to significantly higher levels of cholate compared to untreated communities at the end of antibiotic treatment (Day -2; Figure 3A). Deoxycholate and taurocholate levels were not significantly different between untreated communities and communities treated with antibiotics on Day -2 (Figure 3B-C). There were no significant differences in bile salt levels between untreated and antibiotic-treated communities on Day 0 (Figure 3D-F). There was no correlation between levels of *C. difficile* on Day 7 and cholate levels on Day -2 (Figure 3G) or Day 0 (R^2^=8.5X10^-8^, p=0.997).

**Figure 3:**
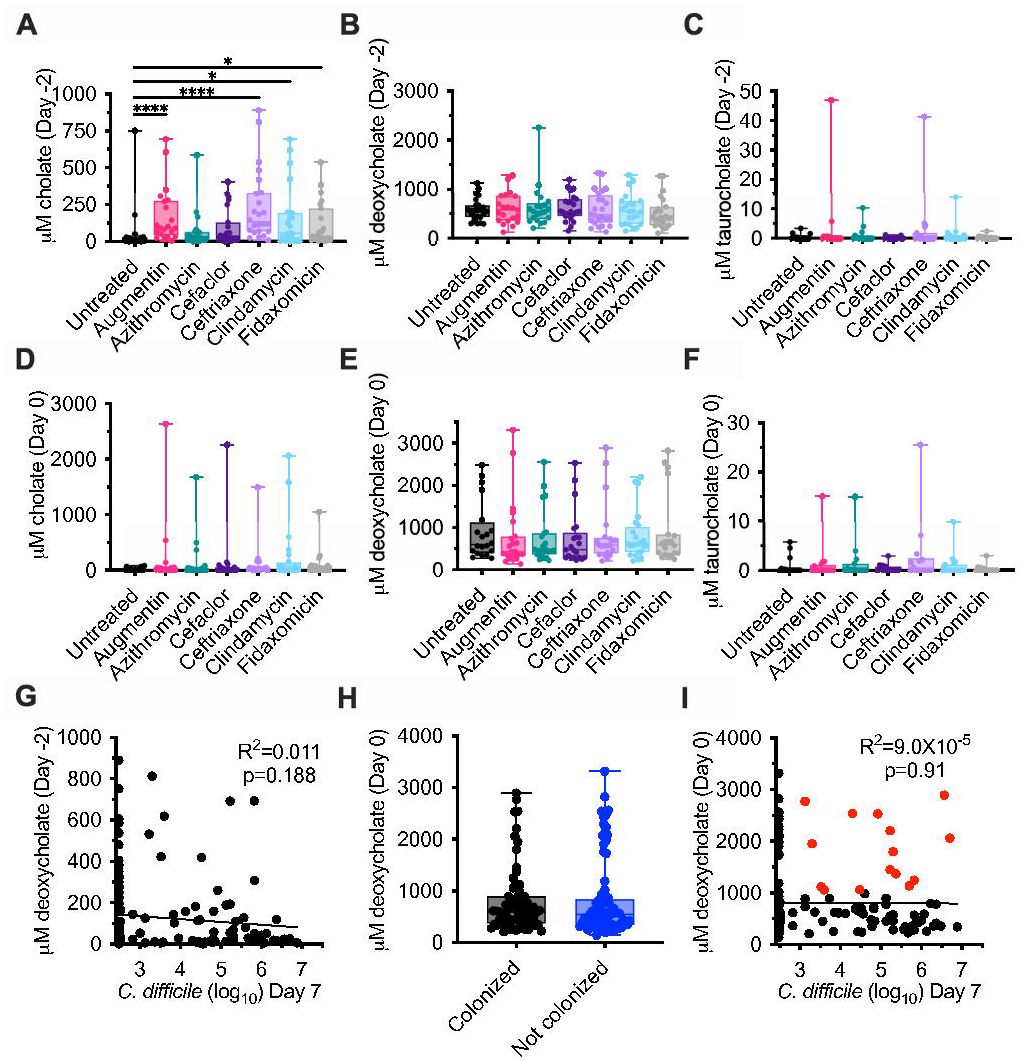
Cholate, deoxycholate, and taurocholate levels in untreated and antibiotic-treated bioreactors. Levels of (**A**, **D**) cholate, (**B**, **E**) deoxycholate, and (**C**, **F**) taurocholate were measured from the communities described in Figure 1 at the end of antibiotic treatment (Day -2; **A**-**C**) and just prior to *C. difficile* challenge (Day 0; **D**-**F**). Significance of differences between antibiotic-treated and untreated samples at each time point are reported. *, p<0.05; **, p<0.01; ***, p<0.001; ****, p<0.0001. (**G**) Simple linear correlation between levels of cholate measured on Day -2 and *C. difficile* levels on Day 7. (**H**) Levels of deoxycholate are plotted for the communities identified as colonized and not colonized in Figure 2. (**I**) Simple linear correlation between levels of deoxycholate measured on Day 0 and *C. difficile* levels on Day 7. Colonized communities with deoxycholate levels greater than 1 mM are indicated in red.

Previous studies have found that deoxycholate inhibits the growth of vegetative *C. difficile* cells (7, 15, 20), with inhibitory concentrations ranging from 0.01 – 0.1% (250 - 2500 μM). The median concentration of deoxycholate measured in untreated communities on Day 0, the day of

*C. difficile* challenge, was 602 μM (Figure 3B); median concentrations in antibiotic-treated communities ranged from 408 μM (Augmentin) to 608 μM (clindamycin), but were not significantly different than those observed in untreated communities (Figure 3E). Because both *C. difficile* susceptibility and deoxycholate levels varied by fecal donor and antibiotic treatment (Figure S7), we tested whether deoxycholate levels on Day 0 correlated with *C. difficile* colonization. We observed that there were no significant differences in levels of deoxycholate between colonized and uncolonized communities (Figure 3H), nor were there significant correlations between deoxycholate levels and *C. difficile* levels on Day 7 (Figure 3I). In addition, we observed that 21.4% of colonized communities (15/70) had deoxycholate levels ≥1000 M on Day 0 (Figure 3I, symbols colored red).

### Bile was not required for *C. difficile* colonization resistance *in vitro*

While the previous results were consistent with the hypothesis that secondary bile salts do not mediate colonization resistance in communities cultured in bioreactors, we could not rule out effects of other bile salts that were not measured. To test whether bile salts were required, we monitored colonization resistance of microbial communities established from five fecal samples cultured in parallel in media with and without added bile. Specifically, replicate bioreactors were established for each fecal sample in media either containing or lacking bovine bile. Triplicate communities were treated with clindamycin as previously described (48), while the remaining three replicates for each type of media were left untreated (Figure 4A). Communities were challenged with vegetative cells of CD2015 and *C. difficile* levels were monitored over time. We found *C. difficile* colonization was similarly suppressed in communities cultured in the presence or absence of bile and that treatment with clindamycin led to similar levels of *C. difficile* colonization (Figure 4B).

**Figure 4.**
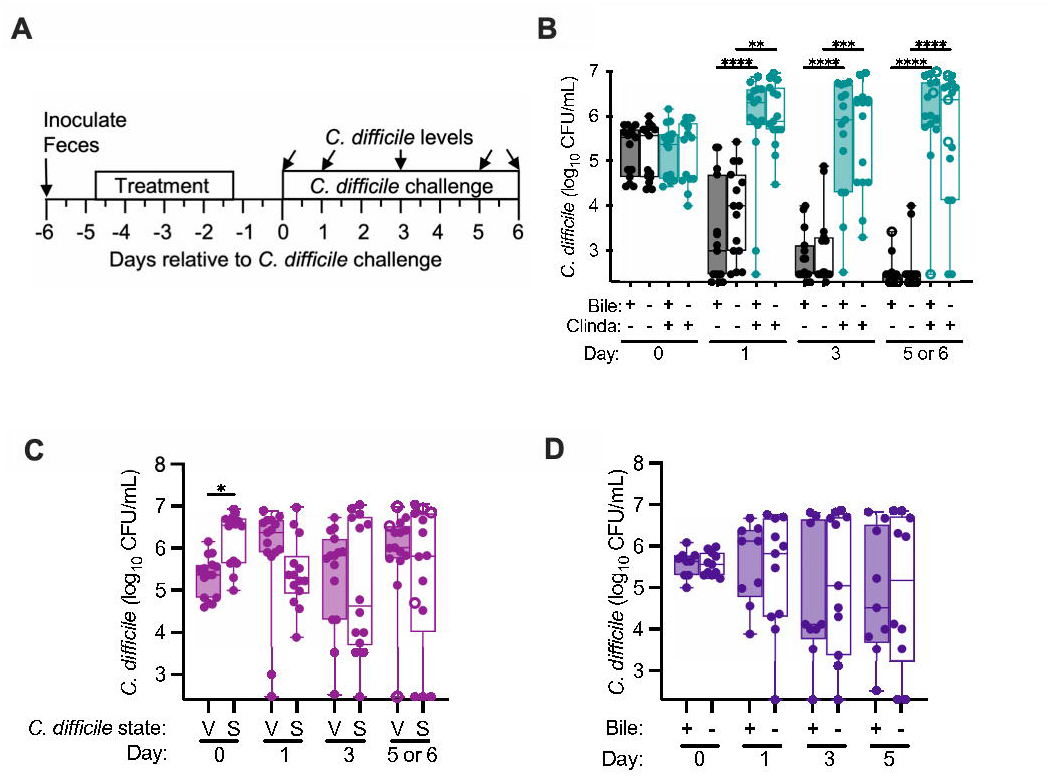
Bile acids were not required for colonization resistance and did not enhance colonization in communities of human fecal microbes cultured in bioreactors. (**A**) Experimental timeline indicating points where samples were collected; time is indicated relative to the point of *C. difficile* challenge. The final collection timepoint was on Day 5 for all samples with the exception of FS22, which was collected on Day 6 and is represented by open symbols (**B**) *C. difficile* levels in untreated and clindamycin-treated communities cultured in the presence and absence of bile. Significance of differences between untreated and clindamycin-treated samples are shown. *, p<0.05; **, p<0.01; ***, p<0.001; ****, p<0.0001. (**C**) *C. difficile* levels in clindamycin-treated communities challenged with *C. difficile* vegetative (V) cells or spores (S). Significance of differences between vegetative and spores are shown at each time point. (**D**) *C. difficile* levels in clindamycin-treated communities cultured in the presence or absence of bile challenged with *C. difficile* spores. No significant differences were observed between communities cultured in the presence or absence of bile.

### Bile was not required for *C. difficile* spores to colonize *in vitro*

Cholate family bile salts are known to greatly enhance *C. difficile* spore germination during *in vitro* culture (15, 49, 50), although they are not required for spore germination in a mouse model of infection (25). Previously, we reported that we were unable to colonize a clindamycin-treated microbial community cultured in an MBRA model with *C. difficile* spores (39). Because these studies were performed using a microbial community formed from pooling twelve fecal donors, we tested the ability of the spores to colonize clindamycin-treated communities established from fecal samples collected from five individuals. We found that the majority of clindamycin-treated communities could be colonized following challenge with *C. difficile* spores (Figure 4C), although communities from one fecal sample were colonized at a lower frequency by spores than by vegetative cells (Figure S8). We then tested whether bile salts were required for *C. difficile* spores to colonize clindamycin-treated fecal microbial communities cultured in MBRAs. We observed that colonization by spores was not significantly different between communities cultured in media with and without bile (Figure 4D).

Because cholate family bile salts were previously reported to enhance germination, we compared the ability of media with and without bile to enhance colony formation of spores in pure culture. Consistent with previous observations (15, 49), incubation of CD2015 spores in bioreactor medium containing bile for one hour enhanced colony formation by germinated *C. difficile* spores by >400-fold compared to incubation in the presence of bioreactor medium without bile (Figure 5). We next tested how spent culture medium collected from clindamycin-treated communities cultured in the presence or absence of bile would impact spore germination. Low levels of spore germination were observed in spent culture medium from fecal communities cultured in the presence and absence of bile (Figure 5). Altogether, this data indicates that the levels of bile salts present in spent culture medium from communities cultured in the presence of bile do not have significant impacts on germination. Further, while the levels of spore germination and colony formation in spent culture medium are low, these low levels of germination would result in ∼10^3^ vegetative cells, a level previously shown to infect antibiotic disrupted communities (39).

**Figure 5.**
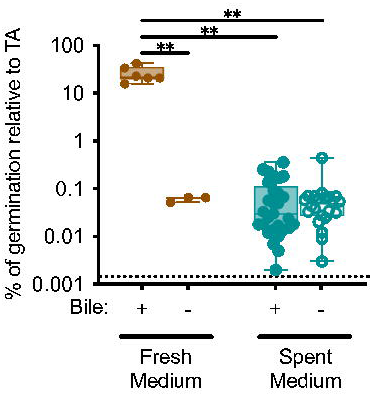
**Low levels of spore germination in spent culture medium from communities cultured in the presence of bile**. Germination of CD2015 spores was determined in filter-sterilized spent culture medium from clindamycin-treated fecal communities cultured in the presence or absence of bile. Spores were incubated in medium for one hour followed by enumeration on agar medium with or without added taurocholate. The percent germination on taurocholate indicates the percent of colonies recovered after overnight incubation in the absence of taurocholate compared to incubation in the presence of taurocholate for each sample. Fresh bioreactor medium made with or without bovine bile were included as controls. Significant difference observed in comparisons between all samples are reported. ns, p>0.05; **, p<0.01; ***, p<0.001.

### Proline metabolism was required for *C. difficile* to colonize a subset of fecal samples *in vitro*

Proteolytic metabolism through Stickland fermentation has previously been reported as an important nutritional niche for *C. difficile* in human (10, 35) and animal models of infection (11, 13, 25, 31, 32, 36, 38). As bioreactor medium contains low levels of fermentable carbohydrates and does not contain mucin-associated monosaccharides, we hypothesized that Stickland fermentation with proline as an electron acceptor could be important for colonization of antibiotic-disrupted fecal communities. To test this hypothesis, we compared the colonization of mutants defective in the ability to metabolize proline (*prdB*::CT (43)) to a wild type (wt) strain; both wild type and *prdB*::CT were in the CD196 background, which is a non-epidemic ribotype 027 isolate (51) . Fecal samples from six healthy individuals were cultured in replicate bioreactors, treated with clindamycin, and challenged with wt or *prdB*::CT strains using the approach described in Fig. 4A.

Overall, we observed that fecal communities challenged with *prdB* mutants showed decreasing levels of colonization over time, with significantly lower levels observed on day 3 and 5 of colonization compared to wild type (Figure 6A). However, the ability of wt and *prdB*::CT strains to colonize was dependent upon the fecal community tested (Figure 6B). Wild-type strains colonized five of the six clindamycin-disrupted communities (FS515, FS228, FS235, FS685, FS133) whereas *prdB*::CT strains failed to colonize two of these susceptible communities (FS228, FS235). In the other three susceptible communities, there was either a lack of *prdB*::CT colonization in the majority of replicates tested (FS515: 10/12 replicates; FS685; 8/13 replicates), or there was no difference between levels of wt and *prdB*::CT colonization levels (FS133). Based on these results, we conclude that Stickland fermentation with proline as an electron acceptor can play an important role in the ability of *C. difficile* to persist in complex communities *in vitro*, but this is dependent upon the composition of the fecal community and its ability to limit *C. difficile* colonization through other mechanisms.

**Figure 6.**
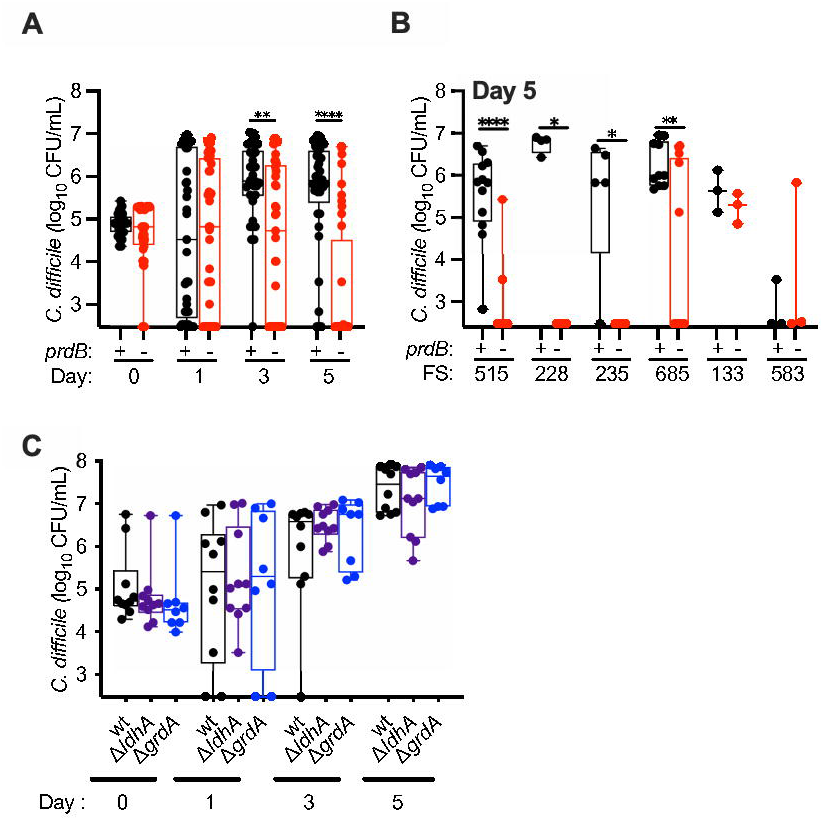
*C. difficile prdB*::CT mutants fail to persist in most clindamycin-treated communities. (**A**) Levels of wt (*prdB*^+^) and *prdB*::CT (*prdB*^-^) CD196 strains were measured over time following introduction into fecal communities treated with clindamycin as outlined in Figure 4A. Data reported are pooled from all six fecal samples tested. Significant differences between wild type and mutant strain at each time point are reported. ***, p<0.001; ***, p<0.0001. (**B**) Levels of wt and *prdB*::CT CD196 strains measured on day 5 of colonization from each of the six fecal samples tested (FS515, FS228, FS235, FS685, FS133, and FS583). Significant differences between wild type and mutant strain for each fecal sample are reported. *, p<0.05; **, p<0.01; ****, p<0.001. (C) Levels of wt, Δ*ldhA*, and Δ*grdA* R20291 strains were measured over time following introduction into fecal communities treated with clindamycin as outlined in Figure 4A. No significant differences between wild type and mutant strains were observed at any time point.

As both glycine and leucine can also serve as electron acceptors for Stickland fermentation, we hypothesized that fermentation of glycine or leucine may also contribute to persistence in clindamycin-treated communities. We tested the ability of mutants defective in Stickland fermentation of glycine and leucine due to mutations in glycine reductase (Δ*grdA*) and 2-hydroxyisocaproate dehydrogenase (Δ*ldhA*), respectively to persist in clindamycin-treated communities. Persistence of these mutants, generated in the hypervirulent ribotype 027 R20291 background, were compared to R20291 wild type strains in four of five fecal samples tested in Figure 6A (FS515, FS228, FS235, and FS133). In these studies, we saw no significant differences in colonization levels between wild type and mutant strains over time, indicating that defects in glycine or leucine fermentation may not be sufficient to limit *C. difficile* colonization in this model.

## DISCUSSION

The factors that govern whether or not an individual exposed to *C. difficile* will go on to develop symptomatic disease are complex. While it is clear that disruption of the GI microbiota is a key risk factor for infection and disease, the extent to which different microbes interact with each other and the host to limit *C. difficile* infection and disease progression are not completely understood. Developing tools that allow microbe and metabolite interactions to be investigated in the absence of a host can help to provide insights into whether mechanisms that inhibit or promote *C. difficile* colonization are more likely to be causative or correlative. This level of understanding is necessary as new therapeutic approaches that more narrowly target *C. difficile*, such as defined microbial consortia are developed.

To gain greater insights into factors that govern *C. difficile* colonization resistance in a fecal MBRA model, we investigated how six different clinically-used antibiotics impacted the ability of the microbiota to resist *C. difficile* colonization. As expected, we observed that antibiotic treatment led to loss of microbial richness, although the extent of microbiota disruption varied both by the class of antibiotic tested and by the composition of the microbial community. Consistent with previous meta-analyses of human antibiotic exposure and risk for *C. difficile* infection in non-hospitalized patients (52, 53), clindamycin was most strongly associated with susceptibility to colonization in antibiotic-treated communities, with ten of twelve communities tested showing susceptibility following treatment. However, the magnitude of microbiota disruption across antibiotic treatments did not correlate with susceptibility to *C. difficile* colonization, with some communities with significant losses in richness resisting *C. difficile* colonization and other communities with modest richness losses exhibiting susceptibility to colonization. This data provides further support for the hypothesis that *C. difficile* colonization resistance is likely due to the presence of specific microbes and their functions that limit *C. difficile* colonization rather than overall microbiota diversity.

Due to the compositional diversity between fecal communities tested, there were relatively few conserved taxa whose presence or absence correlated with susceptibility to *C. difficile* colonization, which is consistent with the model that differences in microbiota function rather than composition are important for understanding *C. difficile* susceptibility. Similar to previous studies (41, 54), however, levels of multiple *Lachnospiraceae* OTUs were higher in resistant communities.

To better understand which microbiota functions may be important for colonization resistance in this model, we investigated two mechanisms previously proposed to regulate *C. difficile* colonization: microbial metabolism of bile acids and Stickland fermentation with proline. glycine, or leucine as an electron acceptor. We observed that microbial communities cultured in bioreactors were able to convert the taurocholate in bovine bile into deoxycholate. Treatment with antibiotics had no significant impacts on levels of taurocholate. Microbes encoding bile salt hydrolases, which remove taurine and glycine from primary bile salts (14), are broadly distributed amongst members of the GI microbiota (55), which may have contributed to the ability of this function to persist during antibiotic treatment.

In contrast, enzymes necessary to perform 7 -dehydroxylation of cholate to deoxycholate are restricted to a smaller subset of the GI microbiota, primarily members of the Clostridiales family and the *Clostridium* genus (56). Treatment with Augmentin, ceftriaxone, clindamycin, and fidaxomicin significantly increased levels of cholate at the end of antibiotic treatment on Day -2, suggesting that microbes capable of 7 -dehydroxylation of cholate to deoxycholate may be lost in some communities, although levels of deoxycholate were not significantly lower at these time points. This lack of significant differences in deoxycholate levels across all fecal sample communities may partly be due to differences in how different fecal communities responded to antibiotic treatment (Figure S7), as some communities (e.g., FS3 and FS9) exhibited much larger decreases in deoxycholate than others (e.g., FS6 and FS12). There was also no correlation between levels of deoxycholate and susceptibility to *C. difficile* colonization, indicating that secondary bile salt mediated inhibition was unlikely to be a mechanism preventing colonization in this model. This was further supported by experiments investigating colonization resistance in the presence and absence of bile, which demonstrated bile was not required for *C. difficile* colonization resistance.

Lack of inhibition by secondary bile salts was surprising, as levels of deoxycholate measured in spent culture medium (431 - 608 μM) were within the range shown to inhibit *C. difficile* growth in pure culture *in vitro* (250 - 2500 μM; (7, 15, 20)). Nevertheless, >20% of communities colonized with *C. difficile* had deoxycholate levels ≥ 1000 M on Day 0 of colonization, suggesting inhibitory effects observed in pure culture may not be the same in complex culture in the presence of other microbes. Further studies are needed to understand whether the presence of other microbes could mitigate some of the toxic effects of deoxycholate on *C. difficile* growth.

We also observed that spores were capable of infecting communities treated with clindamycin and this was independent of the presence of bile. Consistent with what has been previously reported, low levels of germination were observed in the absence of bile and in spent culture medium cultured in the presence or absence of bile. However, these low levels of germination would result in ∼10^3^ vegetative cells, a level previously shown to infect antibiotic disrupted communities cultured in bioreactors (39). This data provides further support to prior studies that have proposed that control of *C. difficile* germination and growth (12, 25) through restoration of microbial bile salt metabolism may not be sufficient to limit *C. difficile* colonization.

While levels of bile salts did not correlate with susceptibility to infection, the ability of *C. difficile* to metabolize proline as an electron acceptor for Stickland fermentation was required for persistence in the majority of disrupted communities tested. These results are consistent with evidence from human (10, 35) and mouse (11, 13, 25, 31, 32, 36) studies that point to Stickland fermentation with proline as an electron acceptor as a preferred nutritional niche for *C. difficile* in the GI tract. Stickland fermentation is a metabolic pathway limited primarily to proteolytic Clostridial species (34). This includes microbes such as *C. scindens*, which can compete with *C. difficile* for proline (25), dehydroxylate primary bile salts to secondary bile salts (24), and can produce a bacteriocin that limits *C. difficile* growth (9). Identifying microbes that effectively compete with *C. difficile* for proline and other amino acids following antibiotic treatment may be one approach to limit *C. difficile* colonization.

It is unclear which metabolites were supporting *C. difficile* growth in antibiotic-treated communities where *prdB* mutants persisted at higher levels over time. Stickland fermentation with glycine or leucine as an electron acceptor is one potential niche that could have been utilized by *prdB* mutants, although we observed the mutants in either pathway alone (Δ*grdA* and Δ*ldhA*) was not sufficient to limit *C. difficile* colonization. Recent studies in a hamster model of infection have shown that hamsters infected with Δ*grdAB* mutants in strain CD630 show delayed morbidity compared to wild type CD630, although there were no significant differences between wt and Δ*grdAB* mutants in levels of *C. difficile* or toxin recovered from the cecum at necropsy (38), indicating the role of glycine fermentation *in vivo* may be relatively modest. Alternatively, *C. difficile* may have utilized carbohydrate fermentation pathways. Further studies are needed to more fully understand how *C. difficile* colonizes disrupted communities.

Altogether, these studies highlight the importance of improving our understanding of bile- independent mechanisms regulating *C. difficile* colonization. While it is clear that Stickland fermentation of amino acids with proline as an electron acceptor is important for *C. difficile* persistence in many disrupted communities, other nutritional niches should also be explored, including whether Stickland fermentation with both glycine and leucine may contribute to reduced persistence. Studies similar to those described here, examining variations in fecal microbial communities and environmental conditions, will further clarify the hierarchical importance of different nutritional environments in limiting *C. difficile* colonization.

## MATERIALS AND METHODS

### Strains used in this study

*C. difficile* strain 2015 (39), a fully-sequenced ribotype 027 isolate resistant to rifampicin and erythromycin (Accession: CP073752.1), was used for routine studies to assess *C. difficile* colonization. To test the role of reductive fermentation of proline in colonization, a previously described ClosTron mutant in proline reductase (*prdB*::ClosTron; (43)) along with its congenic wild type strain were tested. Although originally reported as mutants in the R20291 background, these strains are in the CD196 background (E. Skaar, personal communication). To test the role of reductive fermentation of glycine and leucine, deletions of the coding sequences of glycine reductase (*grdA*) and 2-hydroxyisocaproate dehydrogenase (*ldhA*) were generated by riboswitch-mediated allelic exchange in strain R20291 (57). Successful deletion of genes and absence of second site mutations was verified by Illumina sequencing. Specifically, short read, whole-genome Illumina sequencing was performed by SeqCoast Genomics (Portsmouth, NH USA). Sequencing reads for mutant starins were both compared to the parental *C. difficile* R20291 strain by Geneious Prime’s Geneious alignment algorithm. Sequence variants were then analyzed and both strains were found to be free of mutations outside of the expected gene deletions. Sequences were deposited to NCBI SRA with accession numbers: SRR31632038 and SRR31632038. These mutants were tested along with their congenic wild-type R20291 strain.

### Fecal sample collection and preparation

Fecal samples were collected from 18 healthy individuals who had not been treated with oral antibiotics within the previous six months. Samples were collected from children aged 4-17 (n=3), adults aged 18-65 (n=12), and older adults aged >65 (n=2). Similar numbers of male and female participants agreed to provide samples. However, only donor age categorization (child, adult, older adult) remained linked to specific fecal samples following de-identification. Studies were designed to collect samples across the age span rather than specifically powered to evaluate differences in community composition by age. All adult participants provided consent to participate in the study. Children provided assent to participate along with parental consent to participate. Protocols for collection and use of fecal samples were reviewed and approved by Institutional Review Boards at Baylor College of Medicine (protocol number H-38014) and University of Nebraska-Lincoln (protocol numbers 18585 and 20186).

Samples were self-collected by participants in commode specimen collection containers, sealed in a plastic bag containing a gas pack (BBL GasPak Anaerobe sachet), packed in ice packs, placed in a sealed container, and returned to the lab within 24 hours as previously described (39). Fecal samples were manually homogenized and subdivided under anaerobic conditions (Anaerobic Chamber with 5% H_2_, 5%CO_2_, 90% N_2_ atmosphere) and frozen at -80°C until later resuspended at 25% w/v in reduced phosphate buffered saline (PBS). Fecal suspensions were vortex-mixed for 5 min at ≥ 2500 rpm, centrifuged at 200 X *g* for 5 min to settle large particulates, and the supernatants were either used immediately or amended with 7.5% dimethylsulfoxide and preserved at -80°C until use. Previous work had shown that freezing samples at -80°C did not significantly change the composition of communities cultured in bioreactors (44).

### Bioreactor experiments

MBRAs were assembled and operated under anoxic conditions (5% H_2_, 5% CO_2_, 90% N_2_) at 37 C as described previously (39). MBRAs are strips of six independent continuous flow bioreactors that operate at a 15 mL volume and are continuously stirred with small magnetic stir bars positioned over a stir plate. Each independent reactor in the array contains three ports, one for delivery of fresh medium at a slow continuous rate, one for removal of waste, and a sample port for sampling of medium and bacteria in suspension. Bioreactor medium version 3 (BRM3;(58); Table S1), which simulates nutritional conditions of the distal colon was used for all studies, with the exception of studies testing the effect of bovine bile, when this component was excluded from BRM3. Fecal suspensions from individual donors were inoculated into sterile, anoxic BRM3 medium at 1% w/v final concentration and allowed to grow for 16-24 hr in batch, prior to initiation of continuous flow of medium at 1.875 mL/hr (16 hr retention time for 15 mL bioreactors). Communities were allowed to equilibrate under continuous flow for 1 or 6 days as indicated in figures. Antibiotics were either administered twice daily to individual bioreactors or added directly to source medium as indicated in figures. Research grade antibiotics were obtained from the following sources: Augmentin (5:1 amoxicillin sodium salt to potassium clavulanate; Research Products International); azithromycin dihydrate (Thermo Scientific Chemicals); cefaclor (Thermo Scientific Chemicals); ceftriaxone sodium salt hemiheptahydrate (Thermo Scientific Chemicals); clindamycin phosphate (TCI America); and fidaxomicin (Apexbio Technology, LLC). Antibiotic concentrations used for twice daily dosing were estimated based upon previously published measurements in human feces and/or bile (59–65), although fidaxomicin dosing was limited by its maximal solubility. Clindamycin concentrations used in media were previously described (42). One or two days following cessation of antibiotics as indicated in figure legends, *C. difficile* spores or vegetative cells were administered to reactors at ∼ 1 X 10^5^ CFU/mL (vegetative cells) or 1 X 10^6^ CFU/mL (spores) and levels were enumerated over time. Samples were collected from bioreactors at the time points indicated using a sterile needle and syringe to collect bacteria in suspension. When appropriate, an aliquot of this sample was used to enumerate *C. difficile* by serial dilution and plating on taurocholate cefoxitin cycloserine fructose agar (TCCFA (39), for CD196 and CD196 *prdB*::CT), TCCFA supplemented with 50 μg/mL rifampicin and 20 μg/mL erythromycin (for CD2015), or TCCFA supplemented with 25 μg/mL kanamycin (for R20291, R20291 Δ*grdA*, and R20291 Δ*ldhA*). Sample were centrifuged at ∼3000 X *g* for 5 min to pellet cells. Supernatants were removed, and pellets and supernatants were stored at ≤ -20°C.

### Microbial community analysis by 16S rRNA gene sequencing

DNA was extracted from cell pellets using the BioSprint 96 One-For-All Vet processing kit (Qiagen) according to instructions with the following modifications. Prior to extraction, cells were resuspended in Buffer ASL (Qiagen) and added to sterile deep well plates (Axygen) containing 0.1 mm Zirconia beads (BioSpec Products). Cells were disrupted by bead beating for 2 minutes at 1800 rpm on a FastPrep 96 homogenizer (MP Biomedicals). DNA was amplified in duplicate with Phusion polymerase using barcoded primers 515F and 806R that target the 16S rRNA gene as previously described (41, 44), then sequenced on an Illumina MiSeq using 2 x 250 kits according to manufacturer’s protocol. All sample processing and sequencing was performed by the investigators at the University of Nebraska-Lincoln using equipment shared by members of the Nebraska Food for Health Center. Fastqs were processed by mothur 1.41.3, removing chimeras identified by uchime, mapping sequences against Silva release 132, and clustering OTUs at 99% identity using the OptiClust algorithm (66–68). Mothur 1.48.1 was used to rarefy samples to 6944 sequences, to calculate alpha (observed OTUs, Inverse Simpson) and beta diversity (Jaccard and Bray-Curtis dissimilarity) metrics on rarefied data. Code for sequence processing can be found in Supplementary File 3; Excel worksheets for processing of beta diversity data can be found in Supplementary File 4. An OTU table of rarefied data can be found in Table S2 and compiled data from sequencing analysis can be found in Table S3.

### Bile salt measurements by liquid chromatography-tandem mass spectrometry (LC-MS/MS)

Quantitative analysis of the bile salt levels contained in conditioned-BRM3 culture media sample supernatants was performed by the Texas Children’s Microbiome Center Metabolomics and Proteomics Mass Spectrometry Laboratory (TCMC-MPMSL) using previously published methods(69, 70). Briefly, conditioned-BRM3 growth media samples were removed from the fecal communities cultured in the individual bioreactors using sterile needles and syringes. Samples were added into sterile tubes, and the bacterial cells were pelleted by centrifugation at 3,000 X *g* for 5 min. Clarified supernatants were then sterile filtered using 96- well 0.2 μM polyvinyldifluoride (PVDF) filter plates by centrifugation at 200 X *g* for 5 min. The 96-well, deep well plates used to capture sample filtrates were covered with silicone cap mats and the capped samples were wrapped in parafilm to ensure the fixture of the capmats during shipment. The cell-free conditioned media samples were stored frozen at -80°C pending shipment to the TCMC-MPMSL. The cell-free conditioned media samples were shipped frozen on dry ice, and upon receipt by the TCMC-MPMSL, were stored at -80°C until analysis.

All cell-free conditioned media samples were thawed at ambient temperature on a laboratory benchtop. Once thawed, the entire sample volume was transferred into individual 0.6 or 1.5 mL Eppendorf tubes, and all samples were vortex-mixed using a multi-tube vortexer. Preliminary work indicated that 10-fold and 1,000-fold dilutions were suitable to measure the taurocholic acid (TCA) content, and the cholic acid (CA) and deoxycholic acid (DCA) content, respectively, in each of the cell-free conditioned media samples using a common linear dynamic range of 0.977-1,000 ng/mL for CA, DCA, and TCA – the two different dilution procedures for each metabolite were described previously (70). These sample dilution steps were performed using a working internal standard (WIS) solution prepared in 1:1 methanol: water that contained 250 ng/mL each of D4-CA and D4-DCA as described (70). A 5 μL volume of sample was injected onto the SCIEX QTRAP 6500-based LC-MS/MS system, and bile acid concentrations in the conditioned media samples were back calculated using the regression parameters of the standard curve as described previously (70). Sample data was filtered to remove any with total concentrations of cholate, deoxycholate and taurcholate that were less than 1 microgram/mL. The median level of total bile salts measured was 246 ug/mL; filtering removed 18 of 336 samples. All bile salt data, including the 18 samples that were filtered, can be found in Table S4.

While the levels of TCA, CA and DCA in fresh bioreactor medium was measured at the time cultured samples were tested (69, 70), concentrations were not reported and were lost to follow up. Therefore, we performed a broad screen of the bile acids/salt content of two independently prepared fresh preparations of BRM3-based bioreactor medium (with bovine bile) as well as a batch of BRM3 prepared without bile using another previously reported targeted metabolomics method that is capable of quantitatively measuring 16 bile acids/salts (71). This expanded method can quantify the bile acid/salt content of microbial media samples: CA, glycocholic acid (GCA), TCA, beta-muricholic acid (β-MCA), DCA, glycodeoxycholic acid (GDCA), taurodeoxycholic acid (TDCA), chenodeoxycholic acid (CDCA), glycochenodeoxycholic acid (GCDCA), taurochenodeoxycholic acid (TCDCA), ursodeoxycholic acid (UDCA), glycoursodeoxycholic acid (GUDCA), tauroursodeoxycholic acid (TUDCA), lithocholic acid (LCA), glycolithocholic acid (GLCA), and taurolithocholic acid (TLCA). Briefly, samples were prepared by diluting a 10 µL volume of a thawed cell-free, conditioned BRM3 media sample in a 90 µL volume of a WIS solution that contains deuterated analogs of each of the analytes listed above at concentrations of 250 nM for each prepared in 1:1 methanol: water (71). The samples were prepared directly in glass autosampler vials and the samples were vortex-mixed briefly prior to injecting a 10 µL volume onto the SCIEX QTRAP 6500-based LC-MS/MS system. Bile acid/salt concentrations in the fresh BRM3 medium samples were back calculated using the linear regression parameters of the standard curve as described previously (71). All samples were tested in triplicate. Bile salt data for media can be found in Table S5. These studies confirmed the absence of major forms of bile acids/salts in BRM3 medium made without bile (Table S5).

### Data visualization and statistical analysis

The implementation of the LEfSe (47) algorithim in mothur 1.48.1 was used to identify potential OTUs correlated with *C. difficile* colonization from rarefied data as described in Supplementary File 3. Abundance data for OTUs correlated with *C. difficile* colonization was manually curated from OTU abundance tables in Excel and visualized in GraphPad Prism version 10.2.3. R studio version 2022.07.0+548 running R version 4.2.3 was used to generate the heatmap in Figure S2. Code and dependent software versions used for data analysis can be found in Supplementary File 3. The remaining statistical analyses and visualization was performed using GraphPad Prism. Unless otherwise noted (Figure 5; Figure S8), significance of differences at a single time point between two treatment groups was determined with a Mann-Whitney test and between more than two groups was determined with Kruskal-Wallis testing with Dunn’s correction for multiple comparisons. In Figure 5 and Figure S8, parametric statistics were used to determine significance due to the low number of replicates in one or more samples tested. One-way ANOVA with Brown-Forsythe and Welch correction for unequal variances and Dunnett T3 correction for multiple comparisons was used in Figure 5, whereas a two-tailed student’s t-test with Welch’s correction for unequal variances was used in Figure S8. For analysis of repeated measures shown in Figures 1C, 1D, Figure S1E and Figure S1F, a mixed effects model was used to infer statistical significance with time and reactor as the fixed effects and treatment as the random effect. For CFU/mL data, values below the limit of detection of the assay (333 CFU/mL), were reported as “300” to facilitate plotting.

## Supporting information

Code for sequence analysis

Worksheet for beta diversity analysis

## ACKNOWLEDGEMENTS

We thank Robert Britton (Baylor College of Medicine) for access to data collected in his laboratory by J.M.A that contributed to Figure 4B and analysis to bile salt levels in medium. We thank former University of Nebraska-Lincoln undergraduate students Keegan Schuchart, Yining He, and Austin Johnson for technical assistance. We thank Eric Skaar for providing CD196 and CD196 *prdB*::CT mutants.

J.M.A., X.H., and A.E.J. were responsible for conceptualization and design of the studies. Mutants in *grdA* and *ldhA* were generated and validated by J.N.B. and J.A.S. Data collection was performed by X.H., A.E.J., T.A.A., A.I.L, T.V.T.D., H.C.M., S.J.H., T.D.H., and J.M.A. Analysis was performed by X.H., A.E.J., T.A.A, S.J.H., K.M.H., T.D.H., A.M.H. and J.M.A. J.M.A. and A.M.H acquired funding for the studies. The initial draft of the manuscript was written by A.E.J. and J.M.A., and all authors revised, edited, and approved the final manuscript . The Texas Children’s Hospital Department of Pathology provides salary support to the Texas Children’s Microbiome Center-Metabolomics and Proteomics Mass Spectrometry Laboratory staff, and purchased all of the reagents, the consumables and durable supplies, and the LC– MS/MS equipment described. Sequence analysis was completed utilizing the Holland Computing Center, which receives support from the UNL Office of Research and Economic Development and the Nebraska Research Initiative.

## DATA AVAILABILITY

16S rRNA gene data has been deposited in NCBI’s Sequence Read Archive (SRA) under BioProject ID PRJNA729569. Sequence data from *grdA* and *ldhA* were deposited to NCBI SRA with accession numbers: SRR31632038 and SRR31632038.

## ETHICS APPROVAL

Protocols for collection and use of fecal samples were reviewed and approved by Institutional Review Boards at Baylor College of Medicine (protocol number H-38014) and University of Nebraska-Lincoln (protocol numbers 18585 and 20186).

## Supplementary File 1: Figures S1 to S8 Figure Legends

Figure S1. Changes in microbial richness and diversity across individual reactors in response to time and treatment with antibiotics. (A,C) Microbial richness, (B,D) diversity, and (E,F) community dissimilarity were re-plotted and re-analyzed from Figure 1C-1D to show statistically significant differences between treatments at each time point (A,B) and to show values connected over time across individual reactors (C,D). Note that no communities had been treated with antibiotics at Day -7, but reactors would commence treatment after sampling. For comparisons between fecal inocula and Day -7 samples in A and B, all differences were statistically significant with p<0.0001. In (E) and (F), treatment groups are shown in the same order with abbreviated designations for treatment groups. *, p<0.05; **, p<0.01; ***, p<0.001; ****, p<0.0001.

Figure S2. Distribution of 100 most abundant OTUs across fecal inocula and cultured fecal communities prior to antibiotic treatment (Day -7). Log_4_ OTU abundances are indicated as shaded in fecal inocula (gray boxes) and cultured fecal communities (white boxes) for each fecal sample as indicated. Genus-level classifications of each OTU, organized by phyla, are indicated on the y-axis. Full taxonomy can be found in Table S2.

Figure S3. ***C. difficile* levels vary across microbial communities from different fecal samples treated with antibiotics.** *C. difficile* levels measured from duplicate reactors on (**A**) Day 7 and (**B**) Day 3 were pooled and plotted for each fecal sample (FS1-FS12) and each antibiotic tested as indicated. Circles indicate first replicate sample, while squares indicate second replicate samples. Day 3 data is shown for completeness, although it was not used for determining colonization. Statistical analyses were not performed on each treatment because they were not sufficiently powered. Data is shown to indicate trends in variation observed. Points at the lowest point visible on the y-axis values indicate samples with *C. difficile* levels below the level of detection.

Figure S4. **Correlations between levels of *C. difficile* colonization on Day 7 and changes in microbiota composition.** Simple linear regression was used to investigate relationships between *C. difficile* colonization on Day 7 and changes in microbiota composition. (**A**-**B**) Correlation with richness (Observed OTUs) on (**A**) Day -2 and (**B**) Day 0. (C-D) Correlation with changes in richness from Day -7 to (**C**) Day -2 and (**D**) Day 0. (**E**-**F**) Correlation with changes in microbial diversity (Inverse Simpson Measure) on (**E**) Day -2 and (**F**) Day 0. Goodness of fit (R^2^)and significance of deviation from zero (p-value) are reported.

Figure S5. **Correlations between levels of *C. difficile* colonization on Day 3 and changes in microbiota composition.** Simple linear regression was used to investigate relationships between colonization on Day 3 and changes in microbiota composition. (**A**-**B**) Correlation with richness (Observed OTUs) on (**A**) Day -2 and (**B**) Day 0. (**C**-**D**) Correlation with changes in richness from (**C**) Day -7 to Day -2 and (**D**) Day -7 to Day 0. (**E**-**F**) Correlation with changes microbial diversity (Inverse Simpson) on (**E**) Day -2 and (**F**) Day 0. (**G**-**H**) Correlation with changes in Jaccard dissimilarity before antibiotic treatment (Day -7) and after antibiotic treatment on (**G**) Day -2 or (**H**) Day 0. (**I**-**J**) Correlation with changes in Bray-Curtis dissimilarity before antibiotic treatment (Day - 7) and after antibiotic treatment on (**I**) Day -2 or (**J**) Day 0. Goodness of fit (R^2^)and significance of deviation from zero (p-value) are reported.

Figure S6. **Bile salts in bioreactor medium and cultured bioreactor communities.** (**A**) Cholate and (**B**) chenodeoxycholate/ursodeoxycholate family bile salts were measured in two batches of bioreactor medium. Independent batches are indicated by shape, with three technical replicates tested. In (**A** and **B**), the column labeled “Total” indicates the sum of all (**A**) cholate or (**B**) chenodeoxycholate/ursodeoxycholate family bile salts measured. While the levels of taurocholate, cholate and deoxycholate in fresh bioreactor medium was measured at the time cultured samples were tested (described in references 69 and 70), concentrations were not reported and were lost to follow up. (**C**) Concentrations of taurocholate, cholate and deoxycholate in fresh medium and in untreated communities on Day -2 are shown for comparison. For fresh medium, total indicates total cholate family bile salts (including glycocholate) whereas total indicates the sum of taurocholate, cholate and deoxycholate for spent culture medium. Although glycocholate was not measured in spent culture medium, levels would likely be low based on levels of taurocholate measured.

Figure S7. **Changes in levels of cholate and deoxycholate across fecal communities.** (**A-B**) Levels of cholate and (**C-D**) deoxycholate measured from duplicate reactors on Day -2 (**A,C**) and Day 0 (**B,D**) were pooled and plotted for each fecal sample (FS1-FS12) and each antibiotic tested as indicated. Statistical analyses were not performed on each treatment because they were not sufficiently powered. Data is shown to indicate trends in variation observed. Circles indicate first replicate sample, while squares indicate second replicate samples. (**E)** Cholate and (**F**) deoxycholate levels on Day -2 or Day 0 from all antibiotic-treated reactors from a fecal sample were pooled. Statistical significance of bile salt levels on Day -2 were compared to FS3, which had median levels of cholate that were >2.9 fold higher than all other samples (2.9-46 fold higher across all samples). *, p<0.05; **, p<0.01; ***, p<0.001; ****, p<0.0001.

Figure S8: C*. difficile* spores infect clindamycin-treated microbial communities from different fecal samples similarly to vegetative cells. *C. difficile* levels measured from replicate reactors on Days 0, 1, 3 and 5 or 6 are plotted for each fecal sample (FS11, FS16, FS22, FS27, and FS288) infected with *C. difficile* vegetative cells (circles, filled bars) or spores (squares, open bars). n=3 for all conditions except FS288 challenged with spores, which was n=2. All communities were cultured in media containing bile. *, p<0.05, **, p<0.01, ***, p<0.001.

## Supplementary File 2: Tables S1 to S5

Table S1: BRM3 recipe and instructions

Table S2: OTU abundance data for sequenced samples.

Table S3: Compiled data used for Figure 1-3 and Supplementary Figures S1-S7

Table S4: Filtering of raw bile salt data

Table S5: Raw and processed bile salt data obtained from analysis of two independent batches of BRM3

Supplementary File 3: Mothur and R code for analysis of microbiota data.

Supplementary File 4: Excel worksheets for selection of beta diversity values from mothur sequence analysis output.

Supplementary File 5: Tables S6 to S8

Table S6: Huang.final.abund.opti_mcc.0.01.shared

Table S7: Data for simple.metadata.csv

Table S8: Huang.final.abund.opti_mcc.0.01.cons.subsample.cons.taxonomy

