## Supplementary material for "*Clostridioides difficile* colonization is not mediated by bile salts and utilizes Stickland fermentation of proline in an *in vitro* model": Figures S1 to S8

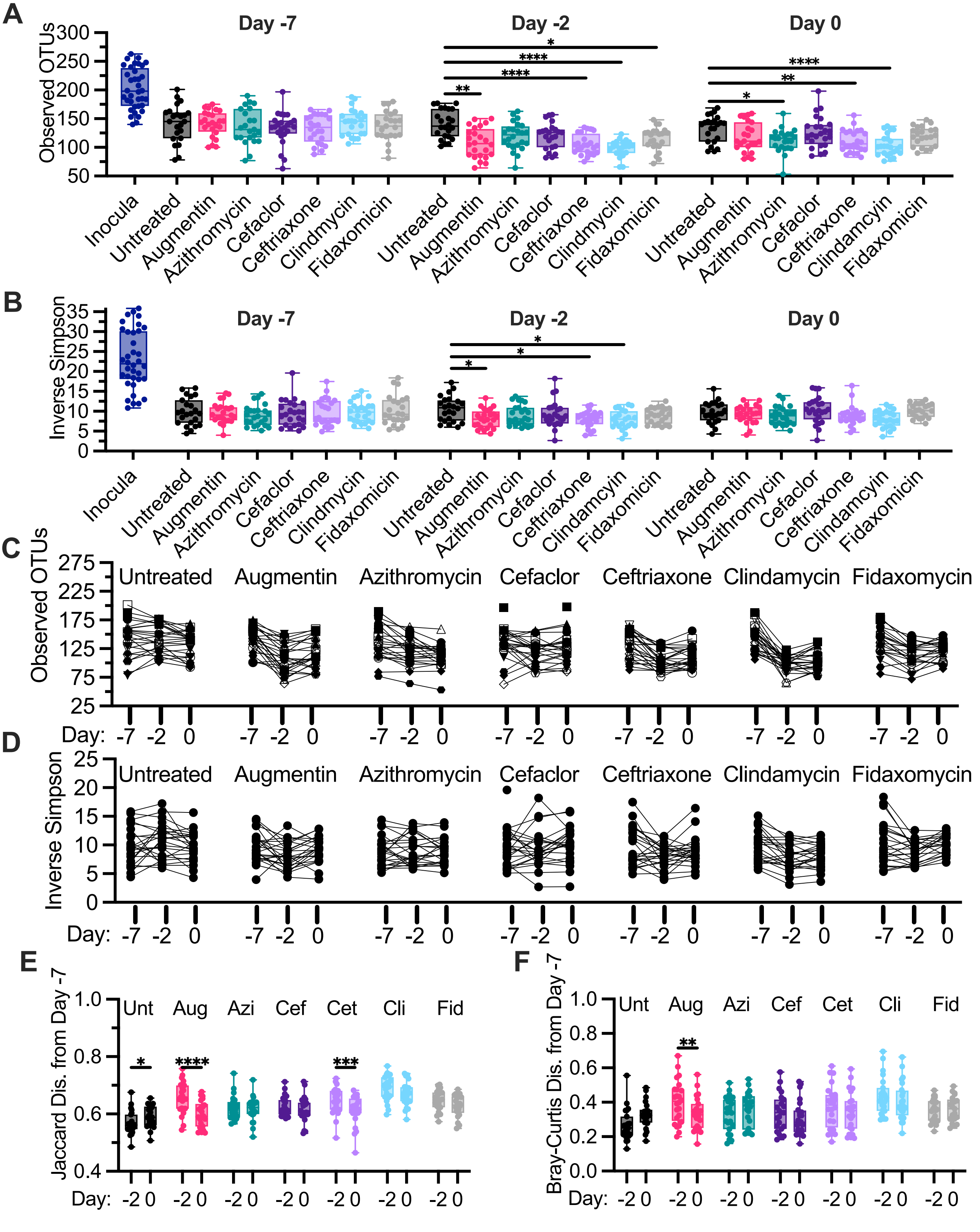


**Figure S1. Changes in microbial richness and diversity across individual reactors in response to time and treatment with antibiotics.** (**A**,**C**) Microbial richness, (**B**,**D**) diversity, and (**E**,**F**) community dissimilarity were re-plotted and re-analyzed from Figure 1C-1D to show statistically significant differences between treatments at each time point (**A**,**B**) and to show values connected over time across individual reactors (**C**,**D**). Note that no communities had been treated with antibiotics at Day -7, but reactors would commence treatment after sampling. For comparisons between fecal inocula and Day -7 samples in A and B, all differences were statistically significant with p<0.0001. In (**E**) and (**F**), treatment groups are shown in the same order with abbreviated designations for treatment groups. *, p<0.05; **, p<0.01; ***, p<0.001; ****, p<0.0001.

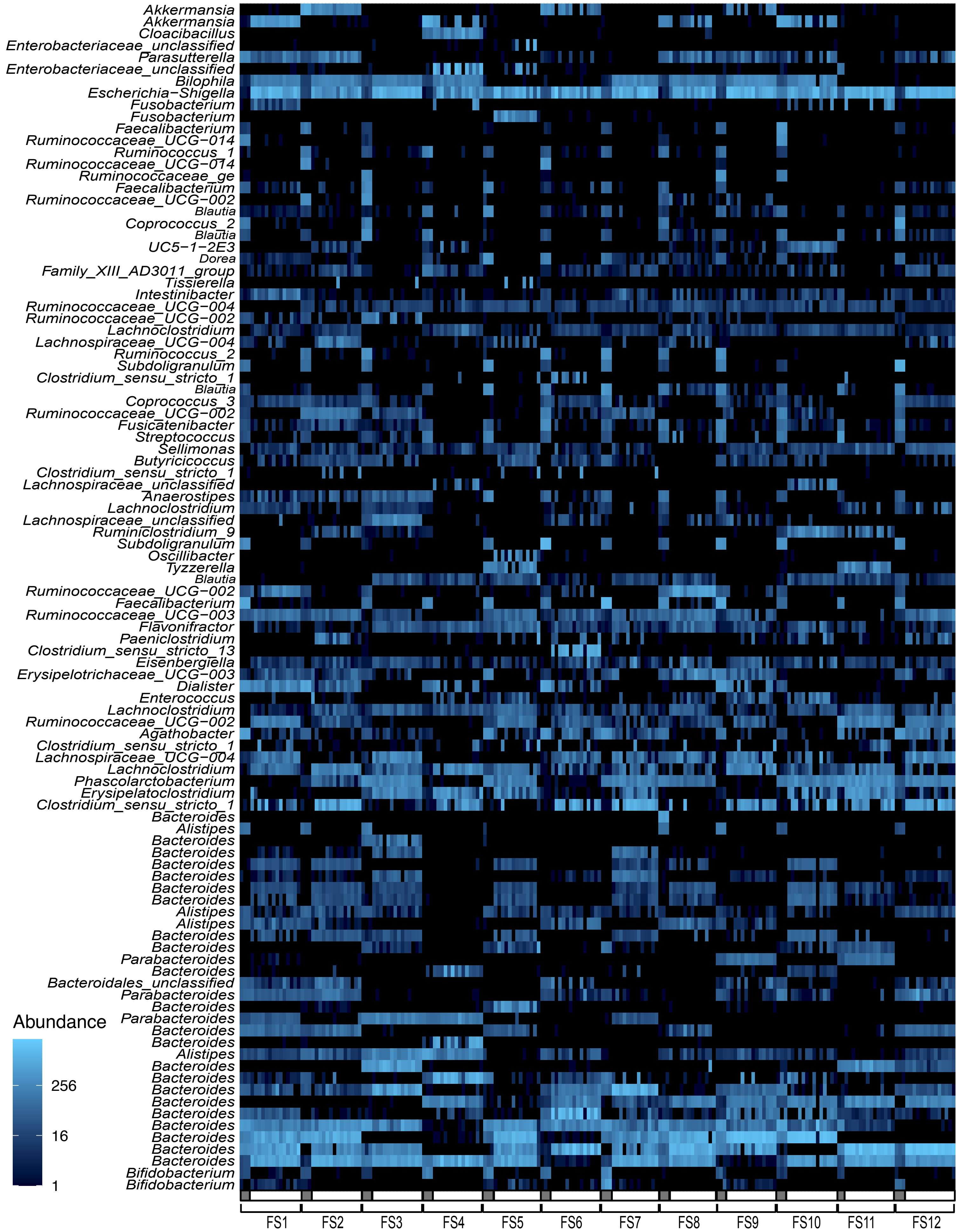


**Figure S2. Distribution of 100 most abundant OTUs across fecal inocula and cultured fecal communities prior to antibiotic treatment (Day -7).** $Log_4$ OTU abundances are indicated as shaded in fecal inocula (gray boxes) and cultured fecal communities (white boxes) for each fecal sample as indicated. Genus-level classifications of each OTU, organized by phyla, are indicated on the y-axis. Full taxonomy can be found in Table S2.

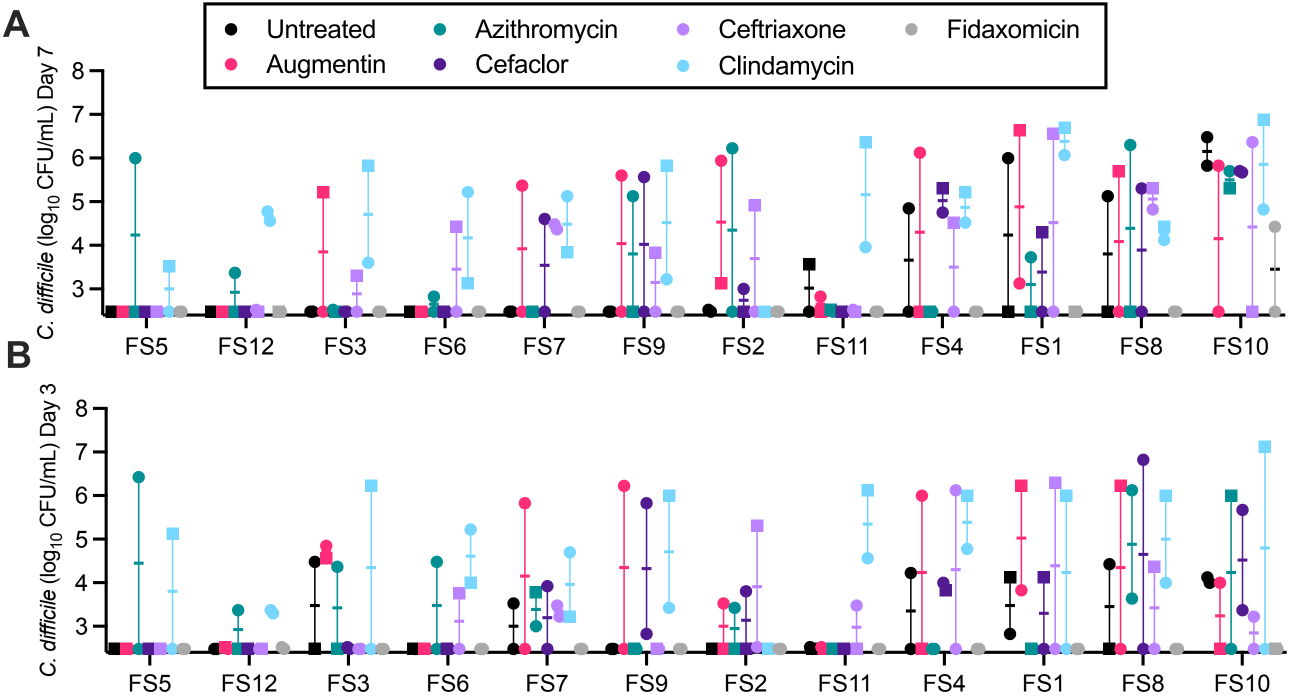


**Figure S3. *C. difficile* levels vary across microbial communities from different fecal samples treated with antibiotics.** *C. difficile* levels measured from duplicate reactors on (**A**) Day 7 and (**B**) Day 3 were pooled and plotted for each fecal sample (FS1-FS12) and each antibiotic tested as indicated. Circles indicate first replicate sample, while squares indicate second replicate samples. Day 3 data is shown for completeness, although it was not used for determining colonization. Statistical analyses were not performed on each treatment because they were not sufficiently powered. Data is shown to indicate trends in variation observed. Points at the lowest point visible on the y-axis values indicate samples with *C. difficile* levels below the level of detection.

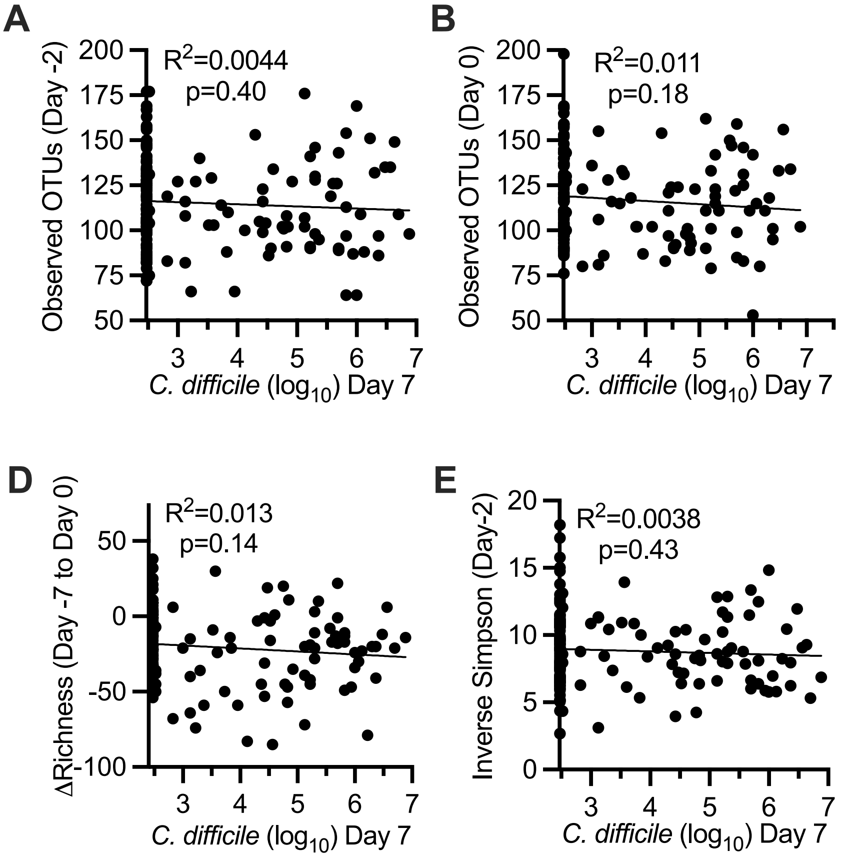


**Figure S4. Correlations between levels of *C. difficile* colonization on Day 7 and changes in microbiota composition.** Simple linear regression was used to investigate relationships between *C. difficile* colonization on Day 7 and changes in microbiota composition. (**A**-**B**) Correlation with richness (Observed OTUs) on (**A**) Day -2 and (**B**) Day 0. (C-D) Correlation with changes in richness from Day -7 to (**C**) Day -2 and (**D**) Day 0. (**E**-**F**) Correlation with changes in microbial diversity (Inverse Simpson Measure) on (**E**) Day -2 and (**F**) Day 0. Goodness of fit ($R^2$)and significance of deviation from zero (p-value) are reported.

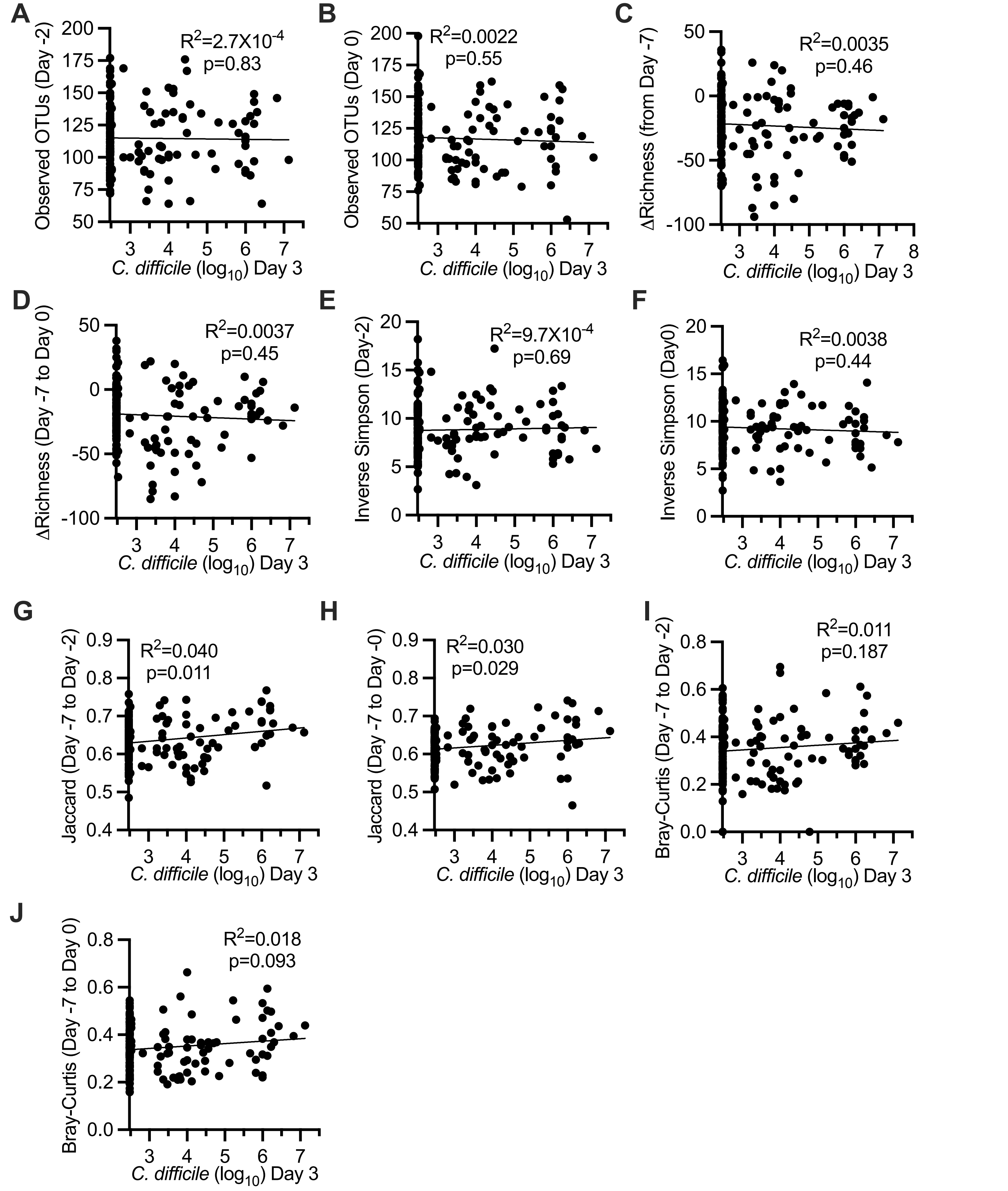


**Figure S5. Correlations between levels of *C. difficile* colonization on Day 3 and changes in microbiota composition.** Simple linear regression was used to investigate relationships between colonization on Day 3 and changes in microbiota composition. (**A**-**B**) Correlation with richness (Observed OTUs) on (**A**) Day -2 and (**B**) Day 0. (**C**-**D**) Correlation with changes in richness from (**C**) Day -7 to Day -2 and (**D**) Day -7 to Day 0. (**E**-**F**) Correlation with changes microbial diversity (Inverse Simpson) on (**E**) Day -2 and (**F**) Day 0. (**G**-**H**) Correlation with changes in Jaccard dissimilarity before antibiotic treatment (Day -7) and after antibiotic treatment on (**G**) Day -2 or (**H**) Day 0. (**I**-**J**) Correlation with changes in Bray-Curtis dissimilarity before antibiotic treatment (Day -7) and after antibiotic treatment on (**I**) Day -2 or (**J**) Day 0. Goodness of fit ($R^2$)and significance of deviation from zero (p-value) are reported.

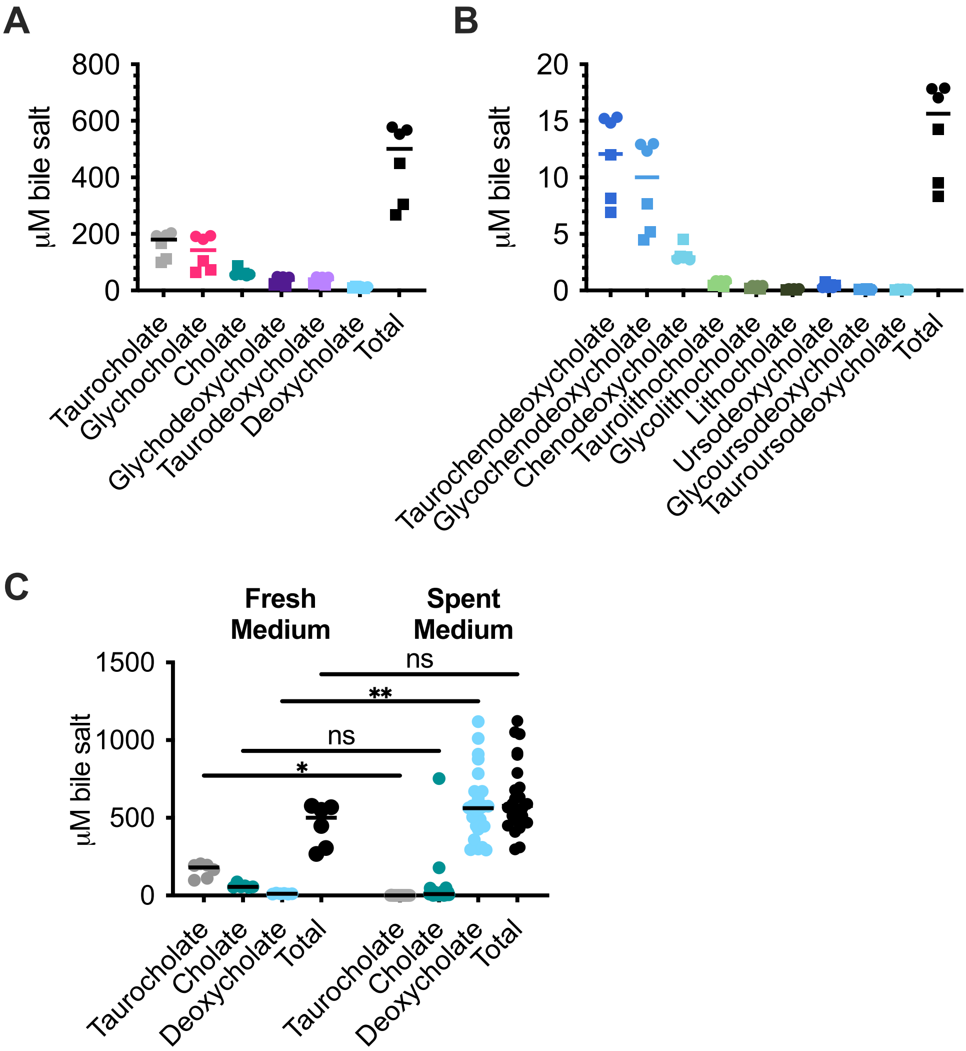


**Figure S6. Bile salts in bioreactor medium and cultured bioreactor communities.** (**A**) Cholate and (**B**) chenodeoxycholate/ursodeoxycholate family bile salts were measured in two batches of bioreactor medium. Independent batches are indicated by shape, with three technical replicates tested. In (**A** and **B**), the column labeled "Total" indicates the sum of all (**A**) cholate or (**B**) chenodeoxycholate/ursodeoxycholate family bile salts measured. While the levels of taurocholate, cholate and deoxycholate in fresh bioreactor medium was measured at the time cultured samples were tested (described in references 69 and 70), concentrations were not reported and were lost to follow up. (**C**) Concentrations of taurocholate, cholate and deoxycholate in fresh medium and in untreated communities on Day -2 are shown for comparison. For fresh medium, total indicates total cholate family bile salts (including glycocholate) whereas total indicates the sum of taurocholate, cholate and deoxycholate for spent culture medium. Although glycocholate was not measured in spent culture medium, levels would likely be low based on levels of taurocholate measured.

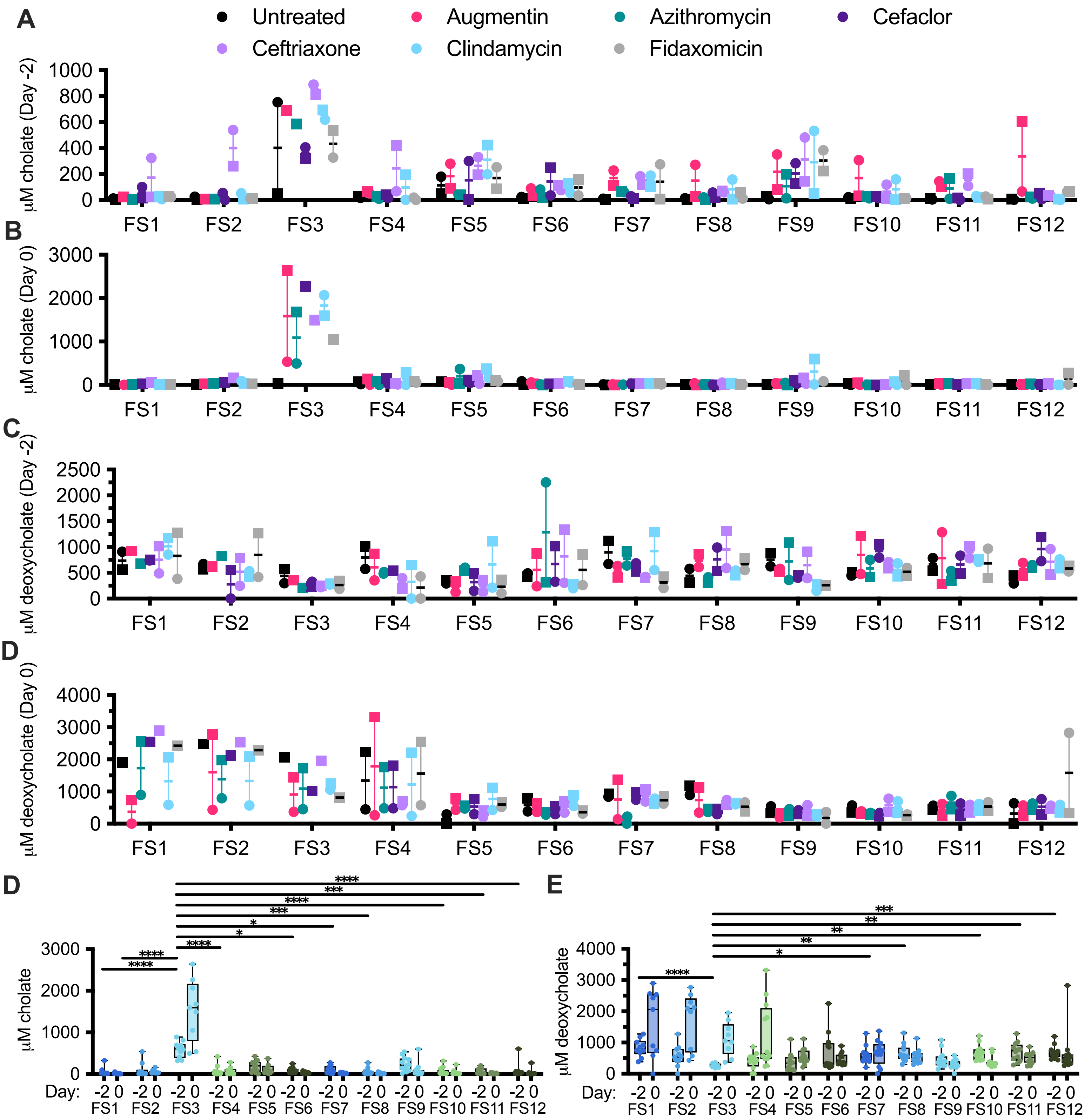


**Figure S7. Changes in levels of cholate and deoxycholate across fecal communities.** (**A-B**) Levels of cholate and (**C-D**) deoxycholate measured from duplicate reactors on Day -2 (**A,C**) and Day 0 (**B,D**) were pooled and plotted for each fecal sample (FS1-FS12) and each antibiotic tested as indicated. Statistical analyses were not performed on each treatment because they were not sufficiently powered. Data is shown to indicate trends in variation observed. Circles indicate first replicate sample, while squares indicate second replicate samples. (**E**) Cholate and (**F**) deoxycholate levels on Day -2 or Day 0 from all antibiotic-treated reactors from a fecal sample were pooled. Statistical significance of bile salt levels on Day -2 were compared to FS3, which had median levels of cholate that were >2.9 fold higher than all other samples (2.9-46 fold higher across all samples). *, $p<0.05$; **, $p<0.01$; ***, $p<0.001$; ****, $p<0.0001$.

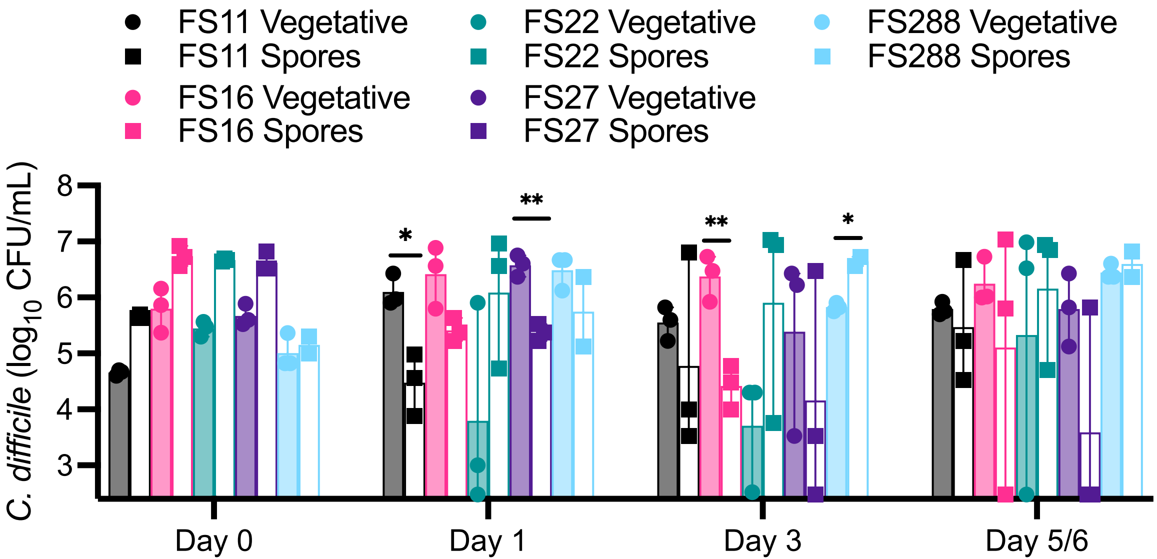


**Figure S8: *C. difficile* spores infect clindamycin-treated microbial communities from different fecal samples similarly to vegetative cells.** *C. difficile* levels measured from replicate reactors on Days 0, 1, 3 and 5 or 6 are plotted for each fecal sample (FS11, FS16, FS22, FS27, and FS288) infected with *C. difficile* vegetative cells (circles, filled bars) or spores (squares, open bars). n=3 for all conditions except FS288 challenged with spores, which was n=2. All communities were cultured in media containing bile. *, $p<0.05$, **, $p<0.01$, ***, $p<0.001$.
